# Evaluating mono and combination therapy of meropenem and amikacin against *Pseudomonas aeruginosa* bacteremia in the Hollow-Fiber Infection Model

**DOI:** 10.1101/2021.12.23.474080

**Authors:** ML Avent, KL McCarthy, FB Sime, S Naicker, AJ Heffernan, SC Wallis, DL Paterson, JA Roberts

## Abstract

Debate continues as to the role of combination antibiotic therapy for the management of *Pseudomonas aeruginosa* infections. We studied extent of bacterial killing and resistance emergence of meropenem and amikacin as monotherapy and as a combination therapy against susceptible and resistant *P. aeruginosa* isolates from bacteremic patients using the dynamic *in vitro* hollow-fiber infection model. Three *P. aeruginosa* isolates (meropenem MICs 0.125, 0.25 & 64 mg/L) were used simulating bacteremia with an initial inoculum ~1×10^5^ CFU/mL and the expected pharmacokinetics of meropenem and amikacin in critically ill patients. For isolates susceptible to amikacin and meropenem (isolates 1 and 2), the rate of bacterial killing was increased with the combination regimen when compared with monotherapy of either antibiotic. Both the combination and meropenem monotherapy were able to sustain bacterial killing throughout the seven-day treatment course, whereas regrowth of bacteria occurred with amikacin monotherapy after 12 hours. For the meropenem-resistant *P. aeruginosa* isolate (isolate 3), only the combination regimen demonstrated bacterial killing. Given that tailored antibiotic regimens can maximize potential synergy against some isolates, future studies should explore the benefit of combination therapy against resistant *P. aeruginosa*.

## Introduction

*Pseudomonas aeruginosa* is the Gram-negative bacteria most commonly associated with mortality and morbidity in hospitalized and immunocompromised individuals (1).

Severe *P. aeruginosa* infections, including bacteremia, require optimised antimicrobial management given mortality rates up to 61% (2). Debate currently exists whether combination therapy can result in better outcomes in the management of severe *P. aeruginosa* infections. Arguments that support the use of combination therapy have done so on the grounds of *in vitro* synergy, an increased likelihood of microbiologically adequate empiric therapy and the potential to prevent development of resistance (3). Observational studies suggest that combination drug regimens are advantageous in those individuals with neutropenia and shock (4, 5). In contrast, meta-analyses performed to date have not demonstrated combination therapy to be more effective compared to monotherapy in the treatment of *P. aeruginosa* infections (6–8).

Aminoglycosides are most often used in combination with β-lactam antibiotics as initial empirical therapy for serious infections, but are not considered appropriate as monotherapy as they have demonstrated inferiority in clinical studies, including higher mortality rates, when compared to β-lactam antibiotics (9–11). These recommendations have been included in the European Committee on Antimicrobial Susceptibility Testing (EUCAST) guidelines (12).

Achieving target pharmacokinetic and pharmacodynamic indices early during the antibiotic course of therapy is associated with an improved patient response and reduced mortality, particularly in serious bloodstream infections (13–15). Wong *et al*. evaluated pooled data from 98 critically ill patients with mono-microbial Gram-negative bacillary bacteremia who were treated with β-lactam antibiotics; the study identified improved rates of clinical outcome as defined by completion of the treatment course without changes to antibiotic therapy, when a free β-lactam antibiotic minimum concentration to minimum inhibitory concentration (*f*C_min_/MIC) ratio greater than 1.3 was achieved (16).

Given the challenges of clinically studying the effect of different antibiotic strategies on patient outcome, useful data can be obtained from *in vitro* models for subsequent informed clinical testing (17). The hollow-fiber infection model (HFIM) can simulate the time course of antibiotic concentrations with a specific elimination half-life at a pre-determined inoculum over clinically relevant durations, both of which are technically difficult with animal *in vivo* models. Results from the HFIM are also well correlated with clinical endpoints for bacterial killing and time course of emergence of resistance (18). Here, we studied meropenem and amikacin as mono- and combination therapy against susceptible and resistant *P. aeruginosa* isolates from bacteremic patients and compared the extent of bacterial killing and suppression of resistance (19).

## Results

### *In vitro* susceptibility and mutant frequency studies

The MICs of meropenem and amikacin for the *P. aeruginosa* isolates are summarized in Table 1. One isolate was resistant to meropenem with an MIC of 64 mg/L (EUCAST breakpoints; susceptible ≤2mg/L, resistant >8 mg/L). The mutant frequency for the *P. aeruginosa* isolates in the presence of amikacin (32 mg/L) was 2.93 x10^-8^ CFU/mL, 3.21 x10^-8^CFU/mL, 8.94 x10^-8^ CFU/mL, respectively for isolates 1 to 3. The mutant frequency for the *P. aeruginosa* isolates grown on meropenem impregnated cation-adjusted Mueller Hinton [(CaMH) agar16 mg/L)] was <4.10 x10^-9^CFU/mL and <5.30 x10^-9^ CFU/mL for isolates 1 and 2, respectively. For isolate 3 the mutant frequency was 7.42 x10^-7^ CFU/mL when grown on meropenem-impregnated CaMH agar (256 mg/L). The amikacin MIC following growth on amikacin-impregnated CaMH agar (64 mg/L) for isolate 1, 2 and 3. The meropenem MIC following exposure to meropenem impregnated agar (fourfold the baseline MIC for each isolate) for isolate 3 was 512 mg/L; whilst no growth was observed for isolates 1 and 2.

**Table 1:** Summary of MICs of meropenem and amikacin for the *P. aeruginosa* isolates.

|  | Meropenem |  | Amikacin |  |
| --- | --- | --- | --- | --- |
|  | MIC | Susceptibility | MIC | Susceptibility |
| Isolate 1 | 0.25 | S | 2 | S |
| Isolate 2 | 0.125 | S | 2 | S |
| Isolate 3 | 64 | R | 4 | S |

S – susceptible; R – resistant

### Pharmacokinetic/pharmacodynamic parameters

The observed meropenem T_>MIC_ was 100% for susceptible isolates (isolate 1 and 2) for both dosing regimens (1 g every 8 hours and 2 g every 8 hours) during the first dosing interval and approximately 3% for the resistant isolate (isolate 3) for the 2 g every 8 hours regimen. The *f*C_max_/MIC of amikacin was between 8.4 and 11.7 for isolates with an MIC of 2 mg/L and between 3.5 and 4.1 for the isolate with an MIC of 4 mg/L.

### Hollow-fiber infection model

For isolates susceptible to both amikacin and meropenem (isolates 1 and 2), monotherapy with either drug rapidly reduced the bacterial density by ~5-log_10_ CFU/mL within 4 hours of initial dosing (Figures 1 and 2). This was comparable with the combination regimens, although the combination achieved the same bacterial killing within 2 hours (Figures 1 and 2). In addition, the combination regimen as well as meropenem monotherapy were able to sustain bacterial killing throughout the seven-day experiment duration. Conversely, amikacin monotherapy was only able to sustain bacterial killing for the first 12 hours of treatment for all isolates (Figures 1 and 2).

**Figure 1:**
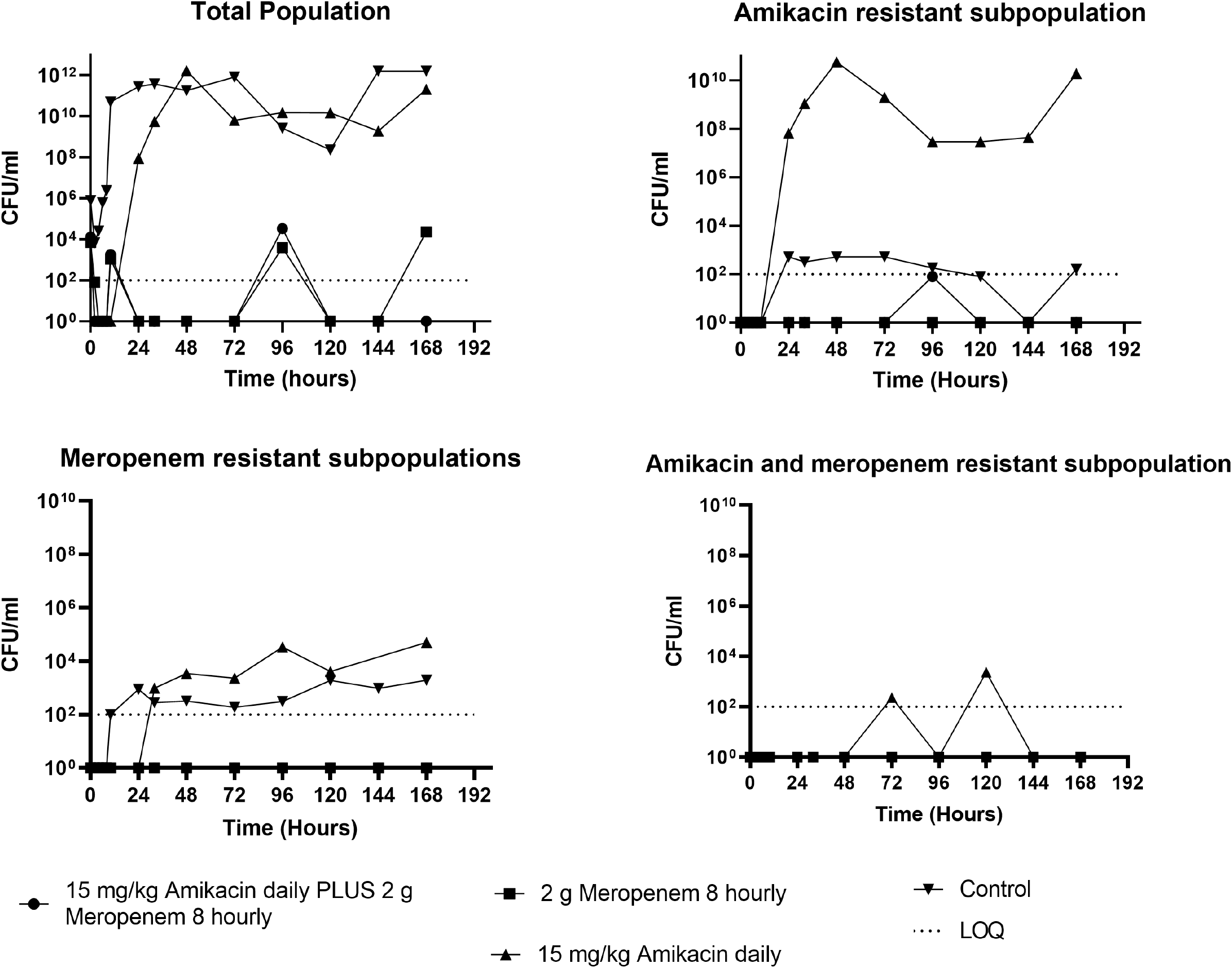
The effect of amikacin or meropenem monotherapy versus amikacin/meropenem combination therapy on the bacterial density of a *P. aeruginosa* isolate (isolate 1: susceptible *P. aeruginosa* isolate) in a hollow-fiber infection model.

**Figure 2:**
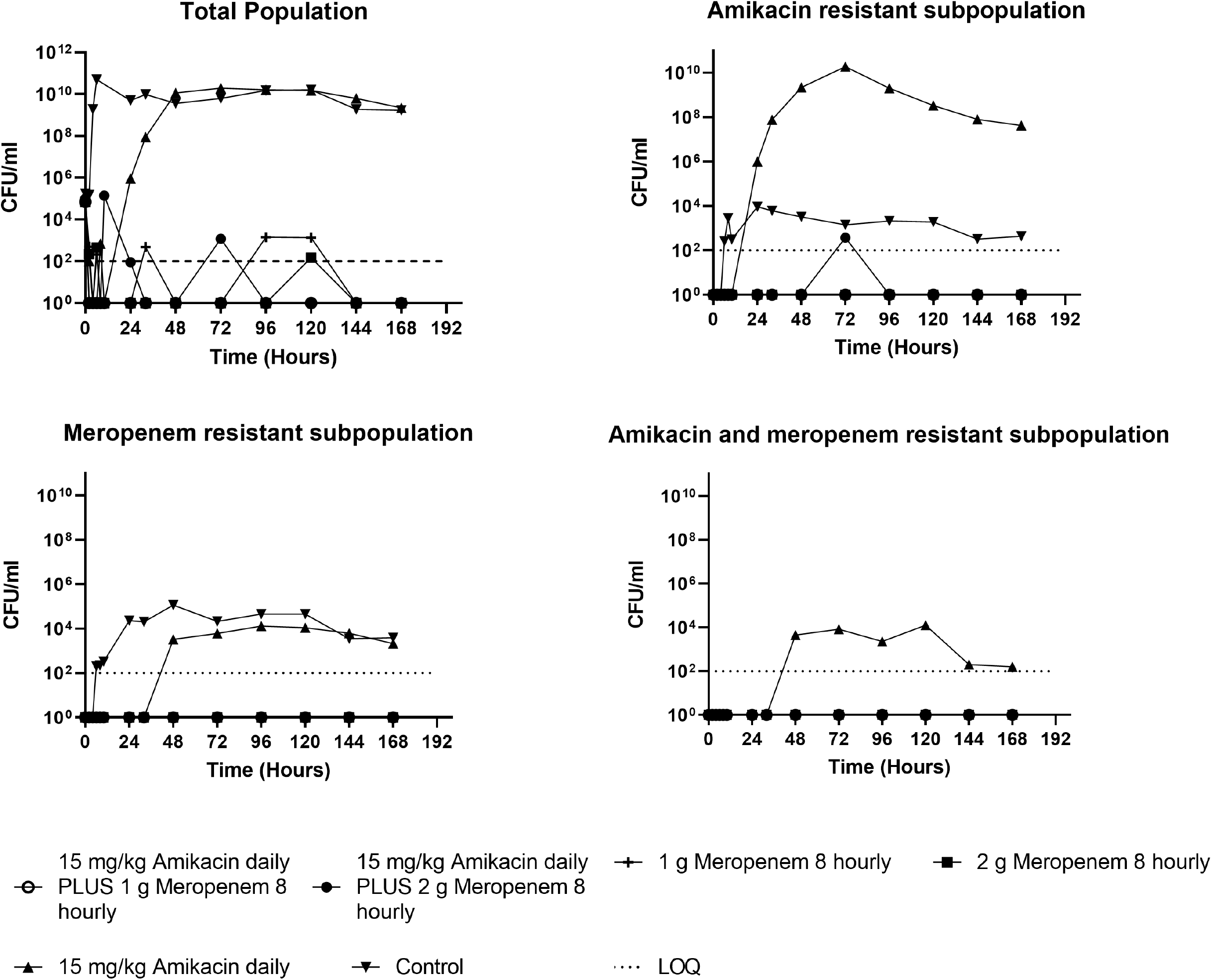
The effect of amikacin or meropenem monotherapy versus amikacin/meropenem combination therapy on the bacterial density of a *P. aeruginosa* isolate (isolate 2: susceptible *P. aeruginosa*) in a hollow-fiber infection model.

Bacterial regrowth on standard CaMH agar mirrored that on amikacin-containing CaMH agar following exposure to amikacin monotherapy (Figures 1 and 2). Resistance to amikacin was confirmed with an MIC increase from 2 mg/L (Table 1) to 64 mg/L and 32 mg/L, respectively for isolates 1 and 2 (Tables 2 and 3). Monotherapy with meropenem did not result in the amplification of growth of subpopulations resistant to either amikacin or meropenem. Treatment with meropenem alone or when combined with amikacin suppressed the growth of a meropenem resistant subpopulation (Figures 1 and 2).

**Table 2:** The MIC of resistant subpopulations emerging during treatment in the hollow-fiber infection model for isolate 1.

| Treatment | Subpopulation | Meropenem MIC (mg/L) | Amikacin MIC (mg/L) |
| --- | --- | --- | --- |
|  |  | Treatment day 7 | Treatment day 7 |
| Amikacin monotherapy | Amikacin resistant | 4 | 64 |
|  | Meropenem resistant | 32 | 64 |
|  | Amikacin and Meropenem resistant | - | - |
| Control (no treatment) | Amikacin resistant | 2 | 64 |
|  | Meropenem resistant | 32 | 64 |

**Table 3:** The MIC of resistant subpopulation emerging during treatment in the hollow-fiber infection model for isolate 2.

| Treatment | Subpopulation | Meropenem MIC (mg/L) |  | Amikacin MIC (mg/L) |  |
| --- | --- | --- | --- | --- | --- |
|  |  | Treatment day 3 | Treatment day 7 | Treatment day 3 | Treatment day 7 |
| Amikacin monotherapy | Amikacin resistant | 0.25 | 0.25 | 32 | 32 |
|  | Meropenem resistant | 16 | 8 | 32 | 32 |
|  | Amikacin and Meropenem resistant | 4 | 8 | 64 | 64 |
| Control (no treatment) | Amikacin resistant | 0.5 | 0.5 | 32 | 32 |
|  | Meropenem resistant | 4 | 8 | 4 | 4 |

For the isolate which was resistant to meropenem (isolate 3), monotherapy with amikacin and combination therapy with amikacin and meropenem reduced the bacterial density ~5-log_10_ CFU/mL within the first 4 hours of treatment (Figure 3). However, monotherapy with amikacin was only able to sustain bacterial killing for the first 8 hours of treatment, whereas the combination sustained bacterial killing for 32 hours after initial dosing. Thereafter, rapid regrowth was observed approximating the initial inoculum within 72 hours (Figure 3 and 4); regrowth mirrored growth on both amikacin and meropenem impregnated CAMH agar. Resistance to amikacin was confirmed with susceptibility testing which showed that the MIC increased from 4 mg/L (Table 1) to 16 mg/L by day seven of treatment (Table 4). The HFIM observed meropenem (Figure 4) and amikacin (Figure 5) concentrations approximated the expected time curve concentrations.

**Figure 3:**
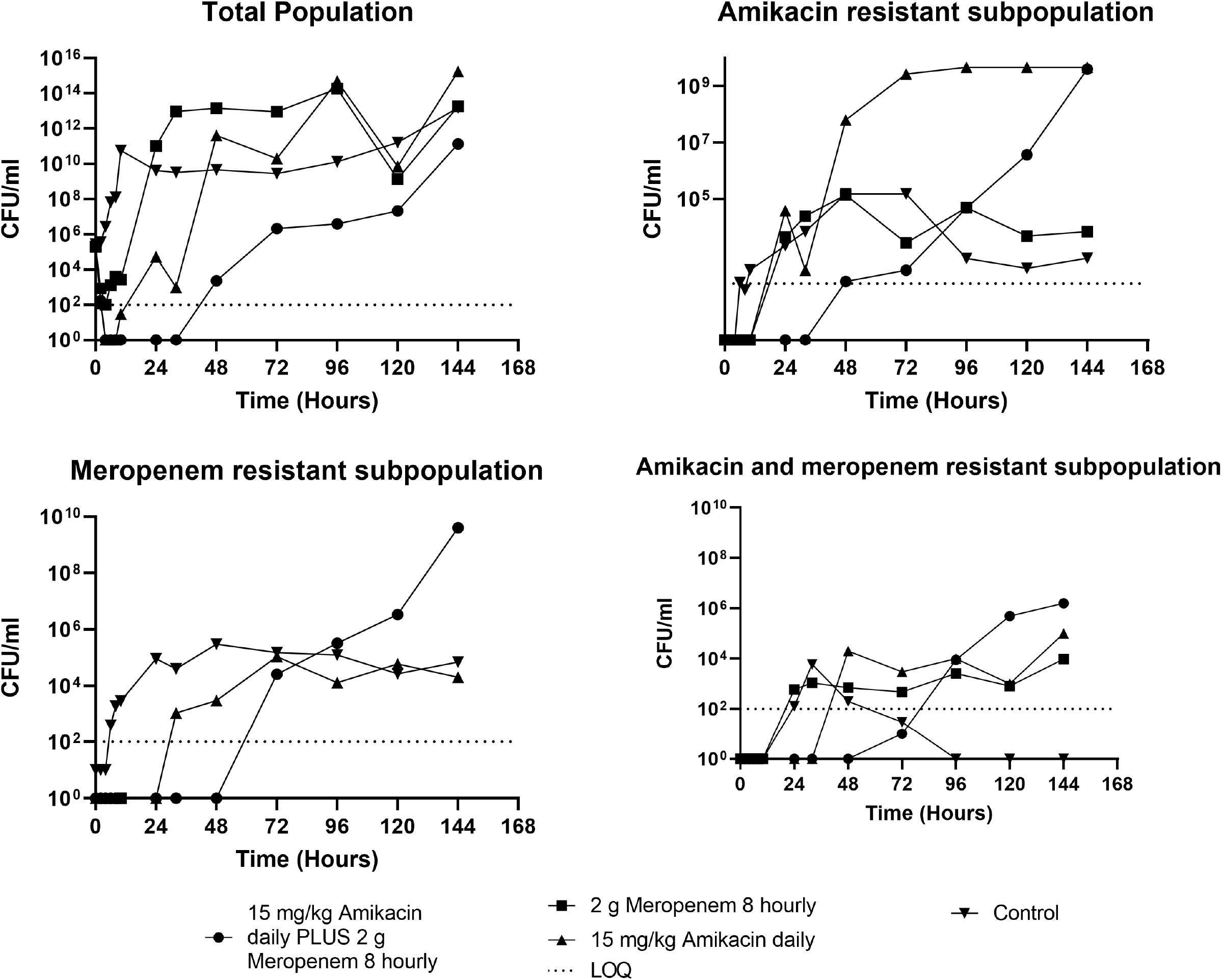
The effect of amikacin or meropenem monotherapy versus amikacin/meropenem combination therapy bacterial density of a *P. aeruginosa* isolate (isolate 3: *P. aeruginosa* isolate resistant to meropenem) in a hollow-fiber infection model.

**Figure 4:**
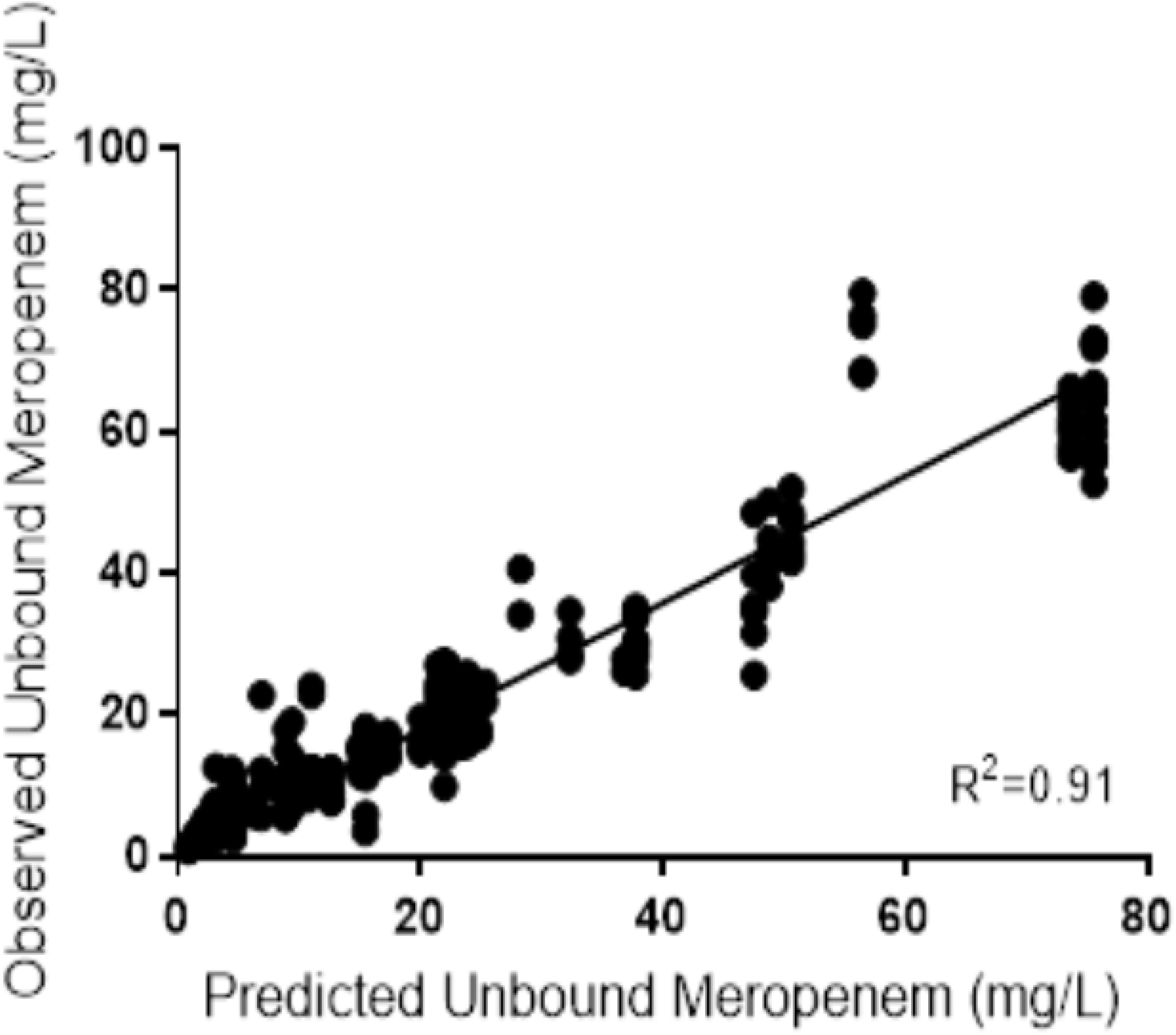
Meropenem observed vs. expected concentration time curve.

**Figure 5:**
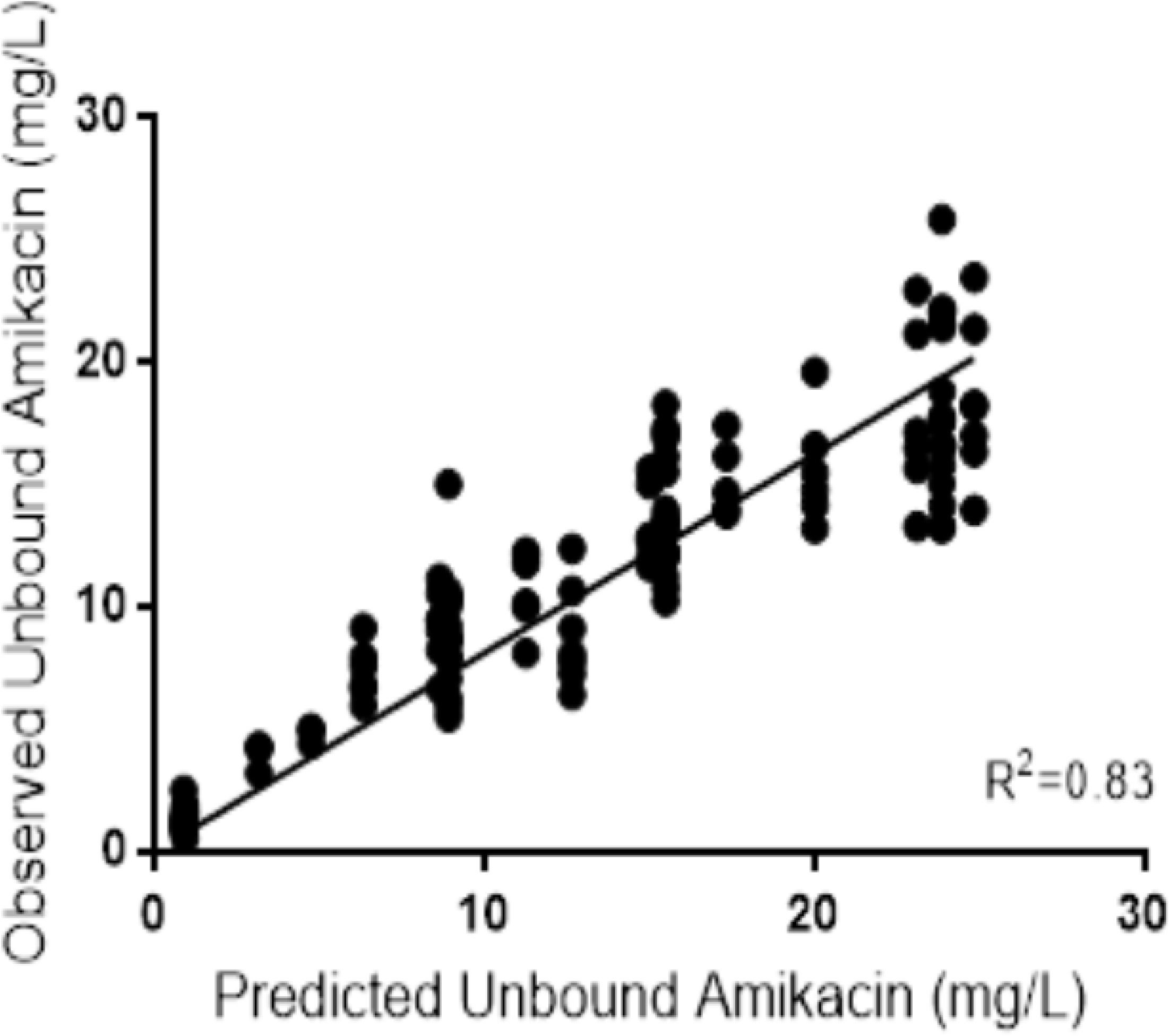
Amikacin observed vs. expected concentration time curve.

**Table 4:** MICs were performed on isolates from each arm and each type of agar plate isolated at the final time point for isolate 3.

| Treatment | Subpopulation | Meropenem MIC (mg/L) | Amikacin MIC (mg/L) |
| --- | --- | --- | --- |
|  |  | Treatment day 7 | Treatment day 7 |
| Amikacin monotherapy | Amikacin resistant | 256 | 16 |
|  | Meropenem resistant | 256 | 4 |
|  | Amikacin and Meropenem resistant | 256 | 16 |
| Meropenem monotherapy | Amikacin resistant | 128 | 16 |
|  | Amikacin and Meropenem resistant | 128 | 16 |
| Amikacin and Meropenem resistant subpopulation | Amikacin resistant | 256 | 32 |
|  | Meropenem resistant | 256 | 16 |
|  | Amikacin and Meropenem resistant | 256 | 16 |
| Control (no treatment) | Amikacin resistant | 256 | 4 |
|  | Meropenem resistant | 256 | 4 |

## Discussion

This study evaluated the impact of meropenem or amikacin monotherapy versus a combination meropenem and amikacin therapy for clinical isolates of *P. aeruginosa* from bacteremic patients. Maximal bacterial killing was similar for both monotherapy and combination therapy isolates susceptible to meropenem and amikacin, although the combination regimen achieved increased the rate of bacterial killing. In addition, the combination regimen as well as meropenem monotherapy were able to sustain bacterial killing throughout the seven-day treatment course whereas regrowth of bacteria occurred with amikacin monotherapy after 12 hours. In contrast, for the *P. aeruginosa* isolate resistant to meropenem (but susceptible to amikacin), only the combination therapy was able to achieve extensive initial bacterial killing, although bacterial regrowth was evident after 32 hours.

Overall, our findings are consistent with that reported previously for meropenem monotherapy and when combined with an aminoglycoside. A previous meropenem HFIM studying the bacterial killing against *P. aeruginosa* in simulated patients receiving 2 g 8-hourly with a creatinine clearance of 120 mL/min also demonstrated a similar bacterial killing profile with sustained bacterial suppression (20). In the previous study, the *f*C_min/_MIC was 2, lower than our own for susceptible isolates with a *f*C_min_/MIC >4. These findings are consistent throughout the literature where a *f*C_min_/MIC between 1 and 4 may suppress regrowth; whereas, a *f*C_min_/MIC >4 commonly suppresses bacterial regrowth (21).

Our findings are also support a previous HFIM study simulating patients with augmented renal clearance receiving combined meropenem and tobramycin (22). Only the continuous infusion meropenem and tobramycin was able to suppress regrowth for 7 days, even at a higher inoculum than what was used in our study (22). This is a common finding in previous time–kill and pharmacokinetic/pharmacodynamic studies (23). The aminoglycoside enhances the beta-lactam antibiotic penetration, thereby increasing the concentration at the target site and associated bacterial killing (24). This would explain the findings of our study whereby the combination of meropenem and amikacin increased the extent and rate of bacterial killing against the meropenem resistant isolate, and even delayed regrowth, when compared with amikacin alone.

Our study also demonstrated regrowth of bacteria following amikacin monotherapy with the emergence of subpopulations resistant to both amikacin and meropenem. This was most likely due to an amikacin resistant subpopulation which existed in the initial inoculum. In addition, a subpopulation resistant to both amikacin and meropenem emerged with amikacin monotherapy, which suggested an amplification of the amikacin resistant subpopulation and potential co-selection of resistance. This has also been described in other HFIM studies where susceptible bacteria were replaced with less-susceptible bacteria following amikacin monotherapy (20). Moreover, Drusano *et al*. described that the resistance mechanism specific for one drug has an impact on the required concentrations of both drugs to suppress the amplification of resistant subpopulations (25), suggesting that optimized amikacin dosing would be beneficial for these patients to reduce the risk of resistance emergence. Although our study did not investigate the potential mechanism underlying the selection of resistance to both amikacin and meropenem following amikacin monotherapy, previous studies have shown aminoglycosides to induce Mex-AB efflux pump expression, leading to carbapenem resistance (26, 27).

The HFIM simulates a patient without an immune system (20). Patients who have the greatest ability to eradicate organisms such as *P. aeruginosa* favor those with a good host immune response, adequate source control and appropriate antibiotic management (28). However, it is challenging to find a model best suited to optimize antibiotic management for those patients who are critically ill and with poor immune functions such as febrile neutropenic patients. Studies examining β-lactam antibiotics, including meropenem, undertaken in the HFIM correlate well with those from neutropenic mouse models and predicts emergence of resistance in patients (29, 30). Drusano *et al*. demonstrated for *P. aeruginosa* in a murine thigh infection that, while granulocytes contribute to the elimination of bacteria up to a certain level, this effect is saturable (31). Moreover, reducing the bacterial density to <1×10^2^ CFU/mL is a likely target for immunocompromised patients to reduce the probability of bacterial regrowth (31). Therefore, our results support the revised EUCAST recommendations of only using aminoglycosides as part of a combination regimen for systemic infections (12, 21). Therefore, patients who are most likely to benefit from combination therapy are those who are immunocompromised, or are likely to have a subtherapeutic β-lactam antibiotic exposure, such as those with augmented renal clearance or those infected with a higher MIC isolate (32).

There are several strengths and limitations associated with our study. We were able to evaluate *P. aeruginosa* isolates that were both susceptible and resistant to meropenem to describe bacterial efficacy for monotherapy or combination therapy. As discussed previously, our HIFM model may be most applicable for an immunocompromised patient; however, characterizing antimicrobial exposures to optimize bacterial killing *in vitro* can aid clinical decision making regarding antibiotic dosing resulting in potentially improved clinical outcomes. We tested conventional doses at only one creatinine clearance; therefore, the bacterial killing and emergence of resistance may be different with a higher or lower simulated antibiotic clearance which would change drug exposures. We only simulated meropenem administered as an intermittent infusion. Previous HFIM studies have shown that the meropenem infusion method has also been shown to be critical for carbapenem-resistant *P. aeruginosa* (22) whereby only a continuous infusion of meropenem when combined with tobramycin suppressed bacterial regrowth. Additionally, we assessed only three isolates and the bacterial killing efficacy and emergence of resistance may differ with other isolates.

The findings of our HFIM study against the tested strains for *P. aeruginosa* isolates support the initial use of combination therapy with meropenem and amikacin in critically ill or immunocompromised patients or in clinical settings where the probability of carbapenem resistance is high. In other clinical scenarios with a *P. aeruginosa* isolate that are susceptible to both meropenem and amikacin, our HFIM supports current meta-analyses that recommend β-lactam antibiotic monotherapy. However, given the strain specific pharmacodynamics of *P. aeruginosa* between the carbapenems and aminoglycosides, it difficult to exclude the possibility that combination therapy may be superior to monotherapy in all situations. Therefore, clinicians may want to consider using the combination therapy for the initial management and ceasing the aminoglycosides once antibiotic susceptibility results have been obtained given the potential nephrotoxicity, vestibular and ototoxicity associated with this class of antibiotics (33).

Combination therapy using both amikacin and meropenem for the initial empiric management of *P. aeruginosa* infections offers some *in vitro* advantages over meropenem monotherapy particularly for immunocompromised patients. Given that optimized dosing of individual antibiotics in combination can maximize potential synergy against some isolates, future studies should explore the conditions for benefit of combination therapy against *P. aeruginosa*.

## Material and Methods

### Antimicrobial Agents

Analytical reference standards of meropenem (as trihydrate, Tokyo chemical Industry Co. LTD) and amikacin (Sigma-Aldrich, PHR1654) were used for *in vitro* susceptibility testing. For dosing simulation in the hollow-fiber infection model experiments, clinical formulation of meropenem (Meropenem trihydrate powder for injection, Ranbaxy Australia Pty Ltd) and amikacin (DBL^™^ Amikacin injection, Hospira Australia Pty Ltd) were used. For meropenem, a fresh 12.5 mg/mL dosing stock solution was prepared from the clinical formulation and kept as 1 mL aliquots in −80°C freezer for preparation of the doses. For each meropenem dose, an aliquot was thawed immediately before dosing and appropriate volume constituted with sterile broth (final volume, 20 mL) into a dosing syringe. Similarly, 10 mg/mL amikacin dosing stock solution was prepared and stored in a refrigerator and appropriate volume constituted with sterile broth (final volume 20 mL) for dosing.

### Bacterial isolates

Clinical isolates of *P. aeruginosa* for three patients were sourced from the University of Queensland Centre for Clinical Research. All bacteria were stored in CaMHCaMH containing 20% glycerol, at −80°C. Prior to each experiment, fresh isolates were grown on Mueller-Hinton agar plates incubated at 37°C for 24h and were used for the preparation of inoculum. For the HFIM studies, the bacterial suspensions for inoculation were prepared form the freshly grown agar plates first by making a 0.5 McFarland standard suspension in sterile water and then subsequently constituting appropriate volume into a 10 mL bacterial suspension in CaMH Broth to achieve a starting inoculum of approximately 1×10^5^CFU/mL to simulate a bloodstream infection. The culture was then incubated at 37°C for 12 hours to give rise to 1×10^9^CFU/ based on a prior growth curve analysis. Finally, an appropriate aliquot of the 12 h culture was diluted with sterile broth (final volume 40 mL) to achieve a final inoculum concentration of approximately 1×10^5^CFU/mL and subsequently confirmed by quantitative cultures.

### *In vitro* susceptibility testing

Broth microdilution method was used for determination MIC of the organism in accordance with the recommendations of the Clinical and Laboratory Standards Institute (CLSI) and EUCAST (34, 35). In brief, serial two-fold dilutions of each antibiotic were prepared in CaMH broth and aliquoted into round bottom microtiter plates (CLSI) and flat bottom microtiter plate (EUCAST). A standardized inoculum suspension prepared in CaMH broth was then added to give a final inoculum of ~5×10^5^ CFU/mL. The inoculated plates were then incubated at 37° C for 16 to 20 hours. The lowest modal concentration of the antibiotic that completely inhibited growth was identified as the MIC in accordance with the CLSI and EUCAST recommendations. The MIC tests were performed on two separate occasions, each with 4 replicates.

### Mutant frequency

A 10-ml culture of a 10^2^ CFU/ml inoculum was incubated in CaMH broth for 24 h at 37°C. Quantitative culturing methods were performed on the resultant bacterial growth using both drug-free CaMH agar plates and antibiotic-containing CaMH agar plates. Antibiotic concentrations were one dilution above the breakpoint for susceptible organisms and two dilutions above the MIC for the resistant organisms. The mutant frequency was taken as the ratio of the concentration of bacterial subpopulations growing on antibiotic-containing plates after incubating for 48 h at 37°C to the total bacterial concentration growing on drug-free agar plates.

### Hollow-fiber infection model

The circuit system for the HFIM was set up as previously described (36). FiberCell Systems cartridge C2011 was used for all experiments. In the experiments investigating the combination of meropenem and amikacin, a supplementing compartment was introduced to simulate the differential clearance of the two antibiotics in accordance with the method described by Blaser *et al*. (37).

The concentration-time profiles of meropenem and amikacin were simulated based on previously described population pharmacokinetic models by Mattioli *et al*.(38) and Romano *et al*. (39) respectively assuming a patient weight of 80 kg and a creatinine clearance of 100 mL/min. This corresponded to simulated half-lives of 1.7 hours and 4.8 hours for meropenem and amikacin respectively. The volume of the central compartment was set at 200 mL and the corresponding systemic clearance calculated for meropenem and amikacin were 1.37 mL/min and 0.48 mL/min respectively. The *f*C_max_ target was 24.8, 42.6 and 85.2 mg/L for amikacin, meropenem 1 g and meropenem 2 g respectively. The *f*AUC target was 171.3, 96.5 and 193.6 mg.h/L for amikacin, meropenem 1 g and meropenem 2 g respectively.

Six separate circuit systems were sets for simulating clinic course of antibiotic therapy for the following regimens: 1) combination therapy with 15 mg/kg amikacin once daily plus 1 g meropenem 8-hourly; 2) combination therapy with 15 mg/kg amikacin daily plus 2 g meropenem 8-hourly; 3) monotherapy with 1 g meropenem 8-hourly; 4) monotherapy with 2 g meropenem 8-hourly; 5) monotherapy with 15 mg/kg amikacin daily; and 6) control (no drug therapy). All doses were administered as a bolus infusion over 30 minutes using a syringe pump and the duration of treatment was seven days. During the treatment period, serial bacterial samples were collected from the extra-capillary space of the hollow-fiber bioreactor before first dose, and at 2, 6, 8, 10, 24, 32, 48, 72, 96, 120, 144, and 168 hours post commencement of treatment. Each bacterial sample were washed twice with sterile phosphate buffered saline. Appropriately diluted bacterial suspension samples were plated on both antibiotic-containing and standard CaMH agar to quantify the likely resistant and total bacterial populations respectively. The drug containing CaMH agar plates included either meropenem, amikacin or the combination of meropenem and amikacin at four-times the baseline isolate MIC.

### Drug assay

Amikacin and meropenem were measured in CaMH broth by a validated chromatographic method. For amikacin analysis 30 μL of a CaMH broth sample was combined with tobramycin (internal standard) and acidified with trichloroacetic acid. An aliquot of the supernatant was injected onto a Nexera2 UHPLC system coupled to an 8030+ triple quadrupole MS detector (Shimadzu, Kyoto, Japan). Chromatographic separation was achieved using a Luna Omega Polar C18 (1.6 μm), 50 x 2.1 mm analytical column (Phenomenex, Torrence, USA) with an ammonium formate / formic acid / acetonitrile mobile phase. Detection of amikacin and tobramycin was performed using an electrospray source in positive mode with optimized multiple reaction monitoring conditions for each analyte. Amikacin was monitored at MRM of m/z 586.25 →163.10 and tobramycin was monitored at m/z 468.20 → 162.95. For meropenem analysis CaMH broth was directly injected onto a Nexera UHPLC system coupled to a Photo Diode Array detector (Shimadzu, Kyoto, Japan). Meropenem was retained away from interferences on a Shim-pack XR-ODS III, 2.0 x 50 mm (1.6 μm) analytical column (Shimadzu, Kyoto, Japan) with a mobile phase of 87% phosphate buffer (0.1 M, pH 7) with 13% methanol. Meropenem was detected at 300 nm. The assay range for amikacin was 1 to 100 μg/mL and for meropenem was 0.5 to 200 μg/mL. The assay methods were validated for linearity, lower limit of quantification, precision, and accuracy according to both U.S. Food and Drug Administration (40) and European Medicines Evaluation Agency criteria. The precision was within 8.9% (amikacin) and 2.2% (meropenem) and the accuracy was within 12.8% (amikacin) and 6.2% (meropenem) at the concentrations tested (amikacin at 1, 3, 10, 40 and 80 μg/mL; meropenem at 0.5, 1.6, 16 and 160 μg/mL).

## ACKNOWLEDGMENTS

The authors would like to acknowledge funding from the Society of Hospital Pharmacists of Australia Merck Sharp and Dohme grant for Pharmacotherapeutics in Infectious Diseases for supporting this research project. Aaron Heffernan would like to acknowledge funding from a Griffith School of Medicine Research Higher degree scholarship. J.A. Roberts would like to acknowledge funding from the Australian National Health and Medical Research Council for a Centre of Research Excellence (APP1099452) and a Practitioner Fellowship (APP1117065).

